# Acquiring Improved Protein Variants With Probabilistic Preferential Learning

**DOI:** 10.64898/2026.06.22.733688

**Authors:** Floris J. van der Flier, Dick de Ridder, Daniel Probst, Henning Redestig

## Abstract

Variant effect prediction (VEP) models can be used to select promising novel enzymes from a pool of candidates. Most supervised VEP models are framed as regression tasks, placing more emphasis on getting the predicted quantities correct than on the relative comparison of individual candidates. Preferential or contrastive models may better align with the goal of selection, or *acquisition*, especially when informed by predictive uncertainty. Here, we introduce a probabilistic preferential learning model based on the Kermut Gaussian process (PKermut) that we designed with the ambition to increase the hit rate among selected variants. We benchmark PKermut against established models, including the original Kermut, the RITA regressor, and an augmented Potts model, on 69 curated ProteinGym datasets across various assay categories.

To evaluate acquisition performance, we propose a novel *quantile* cross-validation scheme that ensures the evaluation of a model’s ability to extrapolate by reserving high-performing variants exclusively for the test set. We assess models using Spearman correlation and evaluate their acquisition performance using five different acquisition functions, encompassing both uncertainty-aware and unaware strategies.

Our experimental results indicate that uncertainty estimates improve the acquisition ability of our models, and that strategies that reward uncertainty generally result in better outcomes than those that do not on single-mutation variant datasets. We observe that PKermut’s Spearman scores and ability to acquire improved variants are greatly affected by the number of variant comparisons sampled in the training set. Kermut achieves the highest Spearman correlation in 54/69 datasets (78%), compared to 12/69 (17%) for PKermut. For acquisition performance, Kermut leads in 44/69 datasets (64%), while PKermut leads in 15/69 (22%). While at this stage PKermut is not a recommended alternative to Kermut, its contrastive nature offers several conceptual opportunities. We share our findings to inspire further development aimed at improving the alignment between training objectives of VEP models and their downstream application in protein engineering.

## 1 Introduction

Enzymes play an essential role in many industrial applications [1]. Protein characteristics specific to the application, like thermal stability, pH tolerance, or surface adsorption, can be strengthened by introducing protein mutations, which has given rise to the field of protein engineering [2]. Success in protein engineering depends on acquisition: the ability to select variants from a pool of candidates that display properties of interest exceeding those of the current best candidate. The more accurately one can pick successful candidates, the fewer engineering cycles are necessary. Nowadays, this identification is predominantly powered by variant effect prediction (VEP) models: computational models that can predict the effects of protein mutations by training on high-throughput screening data. While active learning — iterative cycles of model training and targeted variant screening — has gained increasing attention [3][4][5][6], industrial protein engineering often operates differently. The high-throughput capacity of modern laboratories enables the generation of thousands of variants in a single campaign, and as development progresses, optimization objectives frequently shift once key property thresholds are met, requiring different experimental screening setups. Consequently, batch optimization remains a common paradigm in practice [7]. In this setting, the primary interest lies in exploiting the model’s predictions given a fixed training dataset, rather than adaptively selecting which experiments to run next.

With new VEP models continuously pushing the state-of-the-art [8] on public benchmarks [9], the field of machine learning guided protein engineering is rapidly progressing. However, two related issues are often left unaddressed by novel methodology: (i) a mismatch between performance objectives/assessment and practical use, and (ii) not taking uncertainty in output prediction into account. Underlying the first issue is the fact that the vast majority of published VEP models focus on the prediction of exact variant property values. From the perspective of protein engineering, there is a misalignment between the training objectives of these VEP models and their downstream purpose, which is acquisition. Rather than learning what makes one variant better than the next, the models are trained to accurately predict the properties of every variant in the dataset. The majority of property values are typically distant from the current highest value, and accurately predicting their exact value is of limited relevance to acquisition.

Similarly, practitioners and researchers discriminate between the large body of VEP models by scores that correlate variant property predictions with their ground truth values. Like VEP training objectives, such scores introduce similar misalignments. Ranking scores, like the Spearman correlation reported in benchmarking suites like the ProteinGym [9], serve as a proxy for relative improvement, but conflate meaningful differences among high-performing variants with less relevant comparisons among many low-performing ones, thereby providing an obfuscated measure of the model’s ability to correctly rank high-performing variants. The NDCG score addresses this issue by restricting evaluation to the *k* top-ranked variants. However, for a randomly split dataset, the top *k* variants in the test set generally do not represent the optimal variants across training *and* test partitions. Consequently, the NDCG quantifies ranking performance on an arbitrary subset of variants rather than evaluating the model’s ability to identify the true top performers. While there has been remarkable progress in aligning VEP performance assessment, common scoring metrics are impractical, and no method to evaluate acquisition across models and datasets has been widely adopted yet.

The second issue is related to the typical use of VEP models in protein engineering, *i*.*e*., to guide acquisition by providing predictions that act as a selection criterion, for example, by prioritizing variants whose predicted value exceeds a predefined threshold. However, acquisition based solely on point predictions can be unreliable in the presence of model underspecification [10]. For instance, two candidate sets may have comparable distributions of predicted values, while differing substantially in certain regions of the input space. If one set lies in a well-supported region of the training data and the other in an underspecified region, the resulting acquisition outcomes may differ significantly despite comparable predictions. Uncertainty quantification provides a mechanism for models to reflect this degree of underspecification, enabling acquisition functions to account not only for predicted performance, but also for the reliability of those predictions. While uncertainty quantification is not yet a mainstream feature of VEP models, multiple approaches to uncertainty-guided variant prioritization have emerged in recent years, such as the models proposed by [11] and [8], as well as common adaptations studied by [12]. Kermut stands out among these models, since it reaches state-of-the-art performance on the ProteinGym [8] datasets. Kermut is a Gaussian process that combines sequence and structure representations. However, like most other VEP models, Kermut is trained to perform property value prediction, which may not be to the direct benefit of acquisition. No model with competitive performance on the ProteinGym benchmark currently exists that combines the ability to quantify predictive uncertainty with a contrastive learning objective.

We set out to address these gaps through two separate contributions. First, we develop Preferential Kermut (PKermut), a new probabilistic preferential VEP model. PKermut is a modification of Kermut, equipped with a learning objective centered on correctly predicting the preference relationship between pairs of variants. Second, we present an evaluation framework specifically designed to assess acquisition. Our framework includes a new cross-validation (CV) strategy that creates test folds with domain shift in the property domain, and scores models by their ability to correctly identify high-performing variants in the test fold. We use our acquisition evaluation framework to benchmark PKermut on large number of ProteinGym datasets distributed over different dataset regimes. In addition to comparing with Kermut, we also benchmark two reference VEP models. We evaluate models in terms of the traditional Spearman correlation used for scoring VEP models, and later apply our proposed acquisition-centered evaluation framework to compare pairs of models and acquisition functions. A visual summary of our analysis is given in Figure 1.

**Figure 1:**
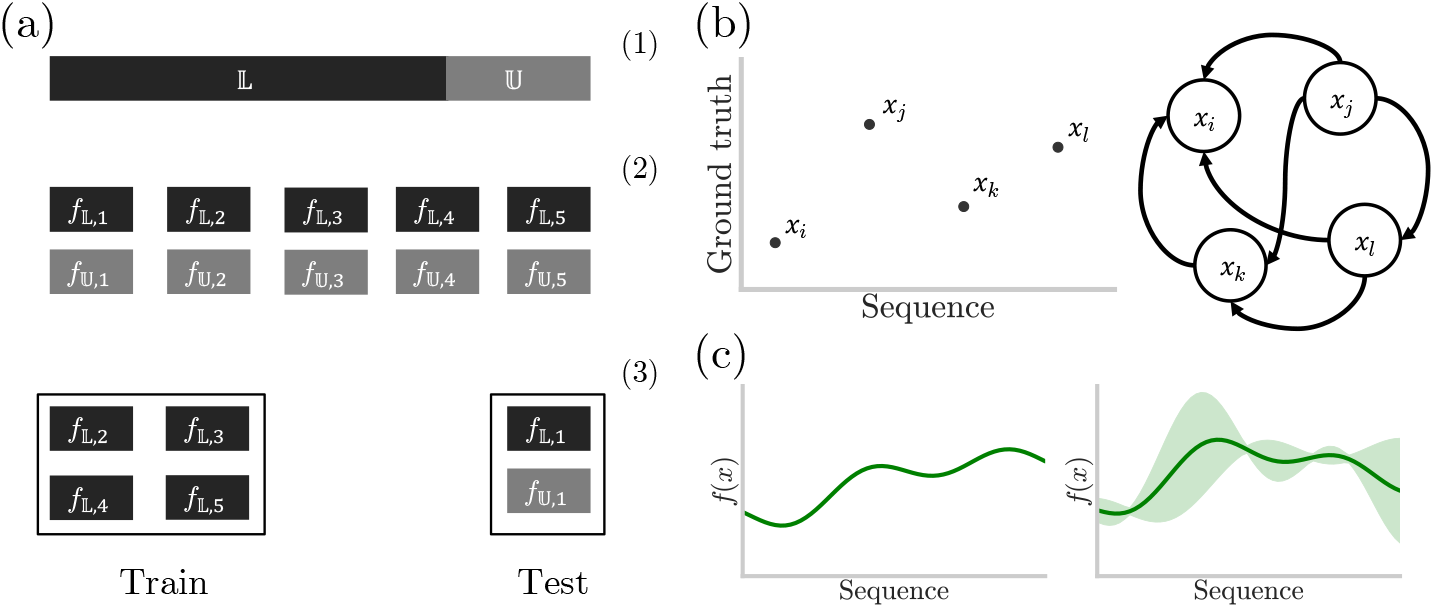
We evaluate how well VEP models prioritize high-fitness variants for experimental testing using 69 ProteinGym datasets, each covering a unique assay category. (a) To assess this ability, we develop the *quantile* cross-validation strategy. This works in conjunction with a new metric termed *recovery* that scores how well a model is able to correctly prioritize top-scoring variants from the test fold. (b) We compare regression-based learning, which predicts exact fitness values (left), against preferential learning, which learns preference relationships (right). The preference structure of a protein variant dataset can be described by a DAG, where directed edges point from higher-valued to lower-valued variants. (c) We evaluate whether uncertainty quantification improves variant prioritization by comparing deterministic VEP models against probabilistic models (state-of-the-art regression and preferential), and comparing uncertainty-aware and uncertainty-unaware acquisition functions across the 69 ProteinGym assays.

## 2 Results

### 2.1 Evaluating acquisition performance of VEP models with and without uncertainty awareness

#### 2.1.1 The *quantile* CV strategy

One way of assessing the acquisition abilities of VEP models in a retrospective fashion, *i*.*e*., using existing data, is to form sets that contain hit variants, with property values exceeding those encountered in the training data, and test how good the models are at prioritizing those variants. Systematically benchmarking models across a wide variety of VEP datasets and comparing acquisition abilities requires a standardized method of data splitting. Such a splitting method should form test sets with predefined ratios of hit variants and ideally should comply with the formation of multiple train and test folds, such that metrics can be collected over non-overlapping test sets and performance metrics can be estimated more reliably than one a single test set, much like standard *k*-fold cross-validation.

Although there have been multiple efforts to systematically assess the extrapolative qualities of VEP models, current methods display limitations in the assessment of acquisition. The *contiguous* and *modulo* splitting strategies proposed by the authors of ProteinGym [9] consider extrapolation in the sequence domain, rather than in the property domain. FLIP introduced the *low-vs-high* split that divides the data into a test set with variants with property values exceeding those of the wild-type, and a training set with values equal to or below that of the wild-type [13][14]. While more closely aligned with the goal of acquiring improved variants, such a split is impractical for datasets produced in later stages of protein engineering development cycles. In such datasets, the property value of the parent protein – the protein from which all variants are derived – is typically much greater than that of the wild-type protein.

To address these limitations, we devised a new cross-validation (CV) scheme based on what we call the *quantile* split. The quantile split divides a dataset into a 75% lower, L, and a 25% upper,U, property interval, the latter containing the 25% best variants with respect to the property of interest. When performing a *k*-fold CV, both quantiles are individually divided into *k* folds. For the *i*^th^ iteration, the training data then contains folds *f*_L,{1,2,…,*k*} *\ i*_, and the test data contains fold *f*_L,*i*_ and fold *f*_U,*i*_. This fold assignment ensures that all hit variants in the test data have property values that exceed those of each variant in the training data. Figure 1(b) illustrates this CV scheme.

#### 2.1.2 Acquiring improved protein variants

We acquire new variants through acquisition functions that use as input a posterior mean and an optional predictive uncertainty produced by any of the models, and output a per-variant acquisition value, *a*, which is used to create a ranking of variants from our acquisition set to prioritize. In the context of batch optimization, a good acquisition function will result in a ranking that prioritizes variants with higher property values in the acquisition set [15]. Acquisition functions can be categorized by whether they account for predictive uncertainty in addition to point predictions. Uncertainty-unaware strategies consider only posterior mean predictions, whereas uncertainty-aware strategies couple those predictions to predictive uncertainty. The latter can be further separated into those that avoid uncertainty, relying on predictions that are best supported by the training data according to the model, and those that favor increased uncertainty, in the hope of finding variants with better properties than the model currently predicts.

Here, we select a range of popular uncertainty-aware acquisition functions reported in earlier works [11, 12] that cover all previously described strategies, and compare them against greedy (*i*.*e*., uncertainty-unaware) acquisition. Specifically, we use upper confidence bound (UCB) and expected improvement (EI) as uncertainty-favoring acquisition strategies, probability of improvement (PI) as an uncertainty-avoiding strategy, and Thompson sampling (TS) as a representative Bayesian optimization strategy commonly used in active learning settings, used to contrast with our batch optimization strategies. We provide more detailed explanations of the acquisition functions included in Methods, section 4.4.

#### 2.1.3 Evaluating acquisition by *recovery*

To systematically evaluate acquisition quality across folds and datasets with the newly described quantile CV strategy, we create a new metric that we term *recovery*, similar to the ‘% of top-100 scores’ metric reported by Greenman et al. [12]. For a given acquisition set and a ranking generated by the acquisition function, recovery is defined as the percentage of hit variants within the first *h* ranked variants in the acquisition set, where *h* is the total number of hit variants in that set. Two important ways in which recovery distinguishes itself from the percentage-based score reported by Greenman et al. are that recovery adapts to the dataset size and is based strictly on variants with property values exceeding those in the training set.

#### 2.1.4 PKermut: preference learning with Kermut

PKermut is a preferential [16] version of the recently released state-of-the-art VEP prediction model, Kermut [8]. Kermut is a Gaussian process trained on variant sequence-property pairs using both sequence and structural features. Unlike standard regression models, PKermut is trained on preference pairs. These are tuples of sample pairs (*u, v*), which indicate that variant *u* is preferred over variant *v*. Importantly, these labels are one-class – only positive preferences are provided. We obtain predictions from PKermut in the form of relative utility predictions per sample, with associated means and variances. For a set of observations, the utility predictions reflect the preference ordering predicted by the model. A more detailed description of PKermut and the underlying pairwise Gaussian process is presented in Methods section 4.2.

To obtain training labels for PKermut, we perform an all-vs-all comparison of property values in our dataset and create a list of preference pairs (*u, v*). The resulting preference structure can be represented by a fully connected directional graph, where the edge *u →v* defines the preference of variant *u* over variant *v*. Without allowing for ties, a preference graph created from *n* variants forms exactly *m* = *n*(*n* − 1)/2 preference edges. Due to the quadratic relationship between *n* and *m*, training on larger datasets could exceed computational constraints. Additionally, since the preference relationship between variants is transitive, the complete graph contains redundancy: we can infer *r →t* if we know that *r → s* and *s→ t*. Thus, we need a method for subsampling *m*, ideally one that maximizes the performance of the model.

Although many such methods can be conceived, in this study, we only consider a uniform preference sampling. One way to reduce the quadratic scaling of *n* with *m* is to subsample the complete preference graph to a smaller preference graph with a predefined verage degree, 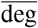. The total number of preference edges in the resulting graph can be calculated by 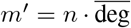. We obtain our subsampled list of preference pairs by uniformly sampling *m*^*′*^ pairs from the complete list of pairs.

### 2.2 PKermut performance is sensitive to preference sampling

<u>We</u> selected three datasets from ProteinGym, displayed in Table 1, to test five values of the subsampling parameter deg and to examine if their impact on PKermut performance varies depending on the size of the dataset. We then train PKermut on all datasets using 5-fold CV with the quantile strategy and calculate the Spearman correlation of the test predictions with the ground truth and the test recovery using greedy acquisition, which we show in Table 1.

**Table 1:** We selected three ProteinGym datasets to perform the pair sampling experiment representing three dataset size categories: Small (*n* = 723), Medium (*n* = 4, 323), and Large (*n* = 9, 462). We then trained PKermut with five uniform pair sampling strategies using 5-fold quantile CV and reported the mean Spearman correlation and recovery with greedy acquisition. Highest scores are displayed in bold and indicate that uniform sampling with 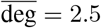 almost consistently results in the best model fit and acquisition.

| Dataset | Category | $\overline{\text{deg}}$ | Spearman ($\rho$) | Recovery (%) |
| --- | --- | --- | --- | --- |
| VG08_BPP22_Tsuboyama_2023_2GP8 | Small | 0.05 | 0.45±0.08 | 38.43±11.31 |
| | | 0.5 | 0.68±0.03 | 40.00±6.69 |
|  |  | 2.5 | **0.77±0.06** | **46.11±8.70** |
| | | 5 | 0.76±0.05 | 43.89±9.09 |
| | | 10 | 0.70±0.08 | 37.22±13.41 |
| HSP82_YEAST_Mishra_2016 | Medium | 0.05 | 0.35±0.08 | 32.50±5.84 |
| | | 0.5 | 0.61±0.03 | 42.04±2.52 |
| | | 2.5 | **0.67±0.03** | 44.72±1.93 |
| | | 5 | 0.67±0.02 | **46.20±3.38** |
| | | 10 | 0.62±0.03 | 41.57±2.25 |
| I6TAH8_I68A0_Doud_515 | Large | 0.05 | 0.20±0.06 | 29.18±5.66 |
| | | 0.5 | 0.49±0.03 | 43.51±2.57 |
|  |  | 2.5 | **0.57±0.05** | **46.00±4.97** |
| | | 5 | 0.43±0.02 | 35.81±1.85 |
| | | 10 | 0.42±0.01 | 34.38±2.46 |

We observe a strong dependency on the number of sampled preference pairs for both model fit and acquisition recovery, with these dependencies showing consistent agreement. In almost all cases, a uniform sampling strategy with 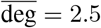 results in both the highest Spearman correlation and greedy acquisition recovery. In subsequent experiments, we therefore train PKermut using a uniform pair sampling with 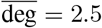.

### 2.3 Model fit comparison on ProteinGym datasets

The ProteinGym contains a large number of VEP datasets with highly variable dataset characteristics, such as the property that is measured, the sequence length, mutational order, etc. However, this space is unevenly sampled, with 64 out of 217 datasets originating from a single study carried out under the same conditions. To obtain a less biased representation of this space, while additionally reducing the number of total training runs, we select a single dataset per assay type, resulting in 66 datasets. Additional criteria used for dataset selection are reported in the Methods, section 4.1.

For our benchmarking experiment, we train PKermut and Kermut with 5-fold quantile CV on all selected datasets. We also include two reference models, the augmented Potts model and the RITA regressor [17], which we describe in Methods section 4.2. While acquisition is our main interest, we first investigate how PKermut compares to other model architectures in terms of the more popular metric, the Spearman correlation. Table 2 lists the average Spearman test scores for every dataset grouping and model, and Figure S1 displays the average Spearman scores for each dataset and model. Although PKermut consistently outperforms the augmented Potts model and the RITA regressor, it is, in turn, marginally outperformed by Kermut in all dataset categories.

**Table 2:** Test-fold averaged Spearman correlation scores and standard deviations for every combination of dataset group and model using the quantile CV strategy. Bold scores display the highest performance by group and indicate that Kermut outperforms all other models in every category.

| Grouping | Group | Count | Model | Speamran ($\rho$) |
| --- | --- | --- | --- | --- |
| All | * | 69 | Kermut | **0.66±0.15** |
|  |  |  | PKermut | 0.62±0.17 |
|  |  |  | Augmented Potts | 0.59±0.15 |
|  |  |  | RITA regressor | 0.57±0.14 |
| Assay type | Activity | 25 | Kermut | **0.60±0.19** |
|  |  |  | PKermut | 0.57±0.20 |
|  |  |  | Augmented Potts | 0.54±0.17 |
|  |  |  | RITA regressor | 0.54±0.17 |
|  | Binding | 8 | Kermut | **0.69±0.14** |
|  |  |  | PKermut | 0.66±0.18 |
|  |  |  | Augmented Potts | 0.61±0.13 |
|  |  |  | RITA regressor | 0.56±0.14 |
|  | Expression | 8 | Kermut | **0.68±0.10** |
|  |  |  | PKermut | 0.64±0.12 |
|  |  |  | Augmented Potts | 0.62±0.09 |
|  |  |  | RITA regressor | 0.58±0.09 |
|  | OrganismalFitness | 26 | Kermut | **0.70±0.11** |
|  |  |  | PKermut | 0.65±0.13 |
|  |  |  | Augmented Potts | 0.62±0.15 |
|  |  |  | RITA regressor | 0.60±0.11 |
|  | Stability | 2 | Kermut | **0.71±0.03** |
|  |  |  | PKermut | 0.69±0.05 |
|  |  |  | Augmented Potts | 0.58±0.05 |
|  |  |  | RITA regressor | 0.58±0.08 |

### 2.3.1 Favoring predictive uncertainty improves acquisition for single mutation variants

Next to scoring the models with the Spearman correlation of their test predictions with ground truth labels, we look at the acquisition recovery. Unlike Kermut and PKermut, the augmented Potts model and RITA regressor cannot leverage uncertainty estimates to inform acquisition. These models are therefore only evaluated with greedy acquisition. Table S1 displays the acquisition recovery statistics for every grouping, model, and acquisition function. Due to the high inter-dataset variance of the recovery, we separately report the percentile ranking of recoveries of model-acquisition function pairs across all datasets in the critical difference diagram in Figure 2.

**Figure 2:**
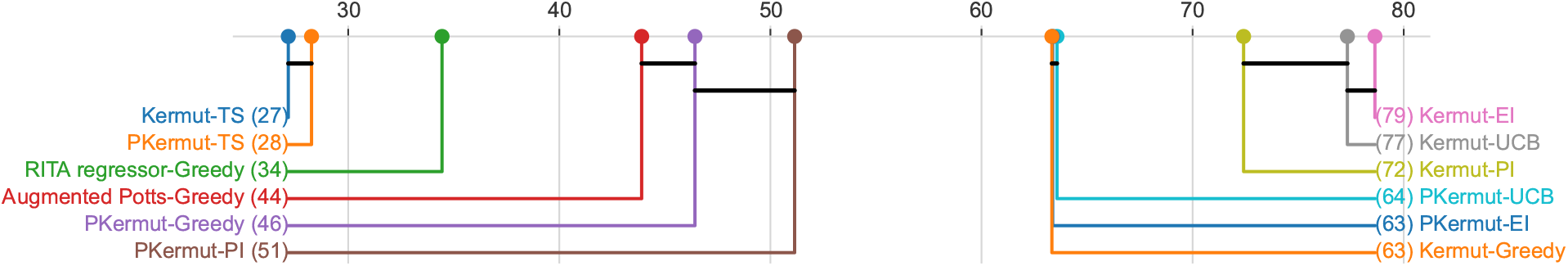
Critical difference (CD) diagram displaying model-acquisition pair strategies, organized from left to right by mean percentile rank of recovery. We ranked the recoveries within each dataset in percentiles and then aggregated the percentile rankings across datasets to obtain a mean percentile rank per strategy. Any two comparisons that are not statistically significant are marked by a horizontal black bar. The results indicate that acquisition with Kermut is favored over all other models, but also that strategies that reward uncertainty are favored over greedy acquisition, and over the probability of improvement, which prefers predictions deemed more reliable by the model.

Again, similar to the Spearman results, we observe that Kermut outperforms all other strategies – *i*.*e*., combination of model and acquisition function – using the EI acquisition function. For both PKermut and Kermut, we observe that acquisition functions that reward uncertainty are significantly favored across all dataset categories. Although these results are limited to single-variant datasets and are obtained in a retrospective setting, they provide evidence that uncertainty estimates can result in improved acquired variants and that these variants reside in regions of increased uncertainty, rather than regions of lower uncertainty. The comparatively poor performance of TS suggests that, within a single iteration of optimization, exploitation-centered batch optimization strategies yield better results than more exploration-centered Bayesian optimization strategies.

## 3 Discussion

We have proposed a new evaluation procedure to benchmark the ability of VEP models to acquire protein variants with improved abilities, an objective central to protein engineering. Our proposed procedure involves a novel CV strategy that creates test folds containing variants with property values exceeding those of the ones in the training folds. The quantile CV strategy works in conjunction with a customized percentage-based acquisition metric that reports the percentage of the total number of hit variants (*h*) found in the test fold within the top *h* ranked variants. We used our acquisition evaluation framework to benchmark PKermut, a modification of the current state-of-the-art VEP model, Kermut. Where Kermut predicts variant property values, PKermut is trained on preference pairs. Both models have the ability to quantify predictive uncertainty, allowing for acquisition with uncertainty-aware functions. Interestingly, our experiments demonstrate that variant acquisition can benefit from selecting variants with increased predictive uncertainty. Not only does this underscore the utility of uncertainty quantification, but it also shows that for specific datasets, improved variants are often found in regions of increased model uncertainty.

### 3.1 Reflections

One of the objectives of this work was to develop a new training routine that is more in line with the goal of selecting promising candidates. We hypothesized that a preferential training objective could offer a more tailored approach by simplifying the task from directly predicting property values to predicting relative improvements. Although Kermut displays a better model fit and ability to acquire improved variants than PKermut in its current form, we believe that PKermut has room for optimization that could result in enhancing its acquisition performance.

Our results indicate that PKermut’s ability to fit the data and acquire improved variants is greatly influenced by the pair sampling strategy, with performance increasing before and tapering off after a certain pair sampling optimum. We hypothesize that performance initially increases due to informative constraints imposed by the preference comparisons. Beyond a certain point, however, the large number of potentially redundant or conflicting constraints may exceed the kernel’s capacity to satisfy them simultaneously, resulting in degraded performance. This is supported by observations where, in certain datasets, adding more comparisons resulted in numerical instability of the kernel matrix. Given that we have tested a small number of naive pair sampling strategies, it is possible that alternative pair sampling strategies result in higher performances. For instance, sampling strategies that assign more weight to pairs of high-valued variants could focus the model on sequence patterns that yield higher property values.

Our second aim was to quantitatively score the acquisition quality of PKermut, Kermut, as well as two reference models, and investigate the use of taking prediction uncertainty into account. Here, we saw that UCB and EI acquisition led to consistently higher recovery scores compared to probability of improvement and greedy acquisition, while Thompson sampling yielded the lowest scores. For single-mutation variant datasets in the ProteinGym, selecting variants with both high posterior mean *and* higher uncertainty predictions is beneficial.

### 3.2 Future directions

The practical value of VEP models extends beyond benchmark metrics to include their ability to adapt to experimental data characteristics. Protein engineering assays are often subject to batch effects, which are left unaccounted for by most models. Preference sampling allows for sampling pairs only (or predominantly) within batches, sacrificing cross-plate preference signal for the elimination of batch effect. The preferences could also be sampled to steer the model to focus its attention on accurately ranking variants in high property domains and ignore irrelevant preferences. Preferential models may also be able to accommodate the multi-objective nature of protein engineering development cycles [18]. Preference labels can be combined across multiple screens to determine a multi-objective preference using a metric like Pareto improvement [19], or a context-dependent preference metric. Aside from preferential training routines, uncertainty-aware acquisition could further be improved by uncertainty calibration [20], parameterizing the acquisition functions, as well as testing other acquisition functions. One could use the quantile CV scheme and the recovery metric as an objective together with search algorithms to find optimized acquisition strategies.

Having demonstrated that uncertainty-rewarding acquisition can lead to better identification of improved protein variants, an interesting next step could be to expand the corpus of public datasets with those displaying characteristics that match protein engineering datasets, and to determine wether uncertainty informed acquisition can lead to increased acquisition on such datasets. This is particularly relevant given the differences between the datasets reported in the ProteinGym and protein engineering datasets. Parent proteins in protein engineering datasets typically have been the result of multiple rounds of optimization, whereas those in the ProteinGym are generally wild types. It can be argued that mutations to naturally occurring proteins more frequently lead to property improvements than to those that have already been specifically engineered to enhance that property. Secondly, all datasets reported here are single-mutation datasets, whereas protein engineering datasets are often of higher mutational order. In the single mutation domain, models are not tasked with learning epistatic relationships between residues.

### 3.3 Conclusion

Our work contributes a new acquisition evaluation framework, which we used to compare a novel preferential model and a number of uncertainty quantification models with the state-of-the-art on a large number of datasets. Our acquisition benchmarking experiment revealed the utility of predictive uncertainty and showed that within the domain of single-mutation protein variant datasets, uncertainty-rewarding acquisition outperforms uncertainty-avoiding acquisition. PKermut represents an attempt at aligning VEP training objectives with protein engineering aims. Although the current implementation did not improve upon Kermut’s performance, we remain convinced that traditional training objectives are poorly suited for downstream acquisition tasks and are optimistic about the potential of preferential learning approaches.

## 4 Methods

### 4.1 Data

We used data from the ProteinGym [9]. To obtain a balanced representation of the assay types and limit computational cost, we selected a single dataset per the “selection assay” category, as included in the metadata. For categories for which more than one dataset exists, we select the one with the highest mutational completeness. We calculate mutational completeness as the number of unique mutations observed across all variants in the dataset, divided by the total possible number of mutations.

We subsequently exclude datasets for which all observed mutations occur at positions spanning less than 10% of all positions in the parent sequence, and also exclude datasets with parent sequences exceeding a length of 1500 residues. Out of the 91 unique selection assay categories, 69 were successfully benchmarked by our analysis. The remaining 22 selection assay categories corresponded to datasets that were either manually excluded or those that exceeded the computational constraints of at least one of our models. We list these categories in Supplementary section A.1.

### 4.2 Models

#### 4.2.1. Glossary

*X* The set of variant sequence observations, *X* = *{x*_*i*_: *i*, …, *n}. u* Variant *u ∈ X*.

*v* Variant *v ∈ X*.

*D* The dataset of *m*^*′*^ preference labels of variant pairs, *D* = *{v*_*k*_ *≻ u*_*k*_: *k* = 1, …, *m*^*′*^*}*.

*f* The latent function generating the variant property values.

***f*** The vector of latent function values.

***θ*** The Gaussian process hyperparameters.

Φ The cumulative density function of a standard Gaussian distribution.

*σ* Noise or scale parameter of the preference likelihood.

Σ The *n × n* covariance matrix produced by the composite kernel function.

Λ The Hessian matrix of the negative log likelihood.

**I** The identity matrix.

### 4.3 Definitions

**Kermut [8]** is a Gaussian process comprising a linear mean function with ESM2 [21] zero-shot predictions. It uses a composite kernel comprising a sequence kernel operating on sequence features derived from ESM2 embeddings, and a structure kernel operating on structure features derived from ProteinMPNN [22].

**PKermut** PKermut shares Kermut’s mean function, composite kernel, and hyperparameter priors, but models preference labels of pairs of variants instead of property values. We implemented PKermut using the PairwiseGP class of the botorch library [23], which based its implementation on the preferential Gaussian process described by Chu et al. [16]. We summarize the core concepts of the preferential Gaussian process in the following paragraphs.

A preferential Gaussian process assumes that the underlying process generating preference labels of any two pairs of variants follows a Bernoulli distribution. We use the notation *u*_*k*_ *≻ v*_*k*_ to indicate that variant *u*_*k*_ is preferred over *v*_*k*_. In an ideal scenario, the likelihood is described by *P*_ideal_:

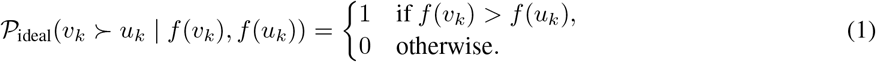

The latent function values of variants are affected by noise, however. Thus, for latent function values *f* (*v*_*k*_), *f* (*u*_*k*_) of variants *u*_*k*_ and *v*_*k*_, assuming Gaussian distributed noise, we can formulate a likelihood as follows:

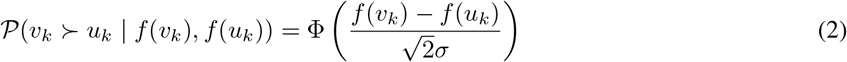

The likelihood over a dataset, *D*, can be formulated as the product of the likelihood function of each constituent preference pair:

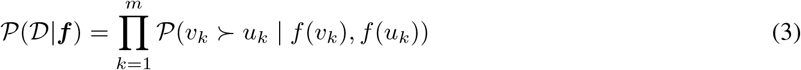

One can parameterize a Gaussian process with a set of hyperparameters, ***θ***, to adapt it to the structure of the data. We can evaluate the likelihood of the data given ***θ*** by the evidence, also called the marginal likelihood:

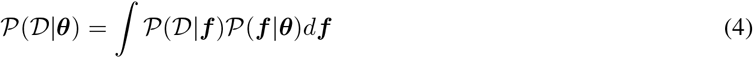

The evidence serves as an objective to find optimal values of ***θ*** (it should be noted that typically its negative logarithm is the loss function that is minimized). In standard Gaussian processes, all terms after the integral are Gaussian, offering an analytical solution for the evidence.

The pairwise likelihood, however, is not Gaussian, meaning that the resulting posterior is also not Gaussian. One can therefore not obtain an analytical solution for the evidence. Chu et al. deal with this limitation by estimating the maximum *a posteriori* function values ***f***_MAP_, and subsequently using Laplace approximation around ***f***_MAP_ to approximate the posterior as a Gaussian. ***f***_MAP_ is found by minimizing the negative log posterior, *S*(***f***):

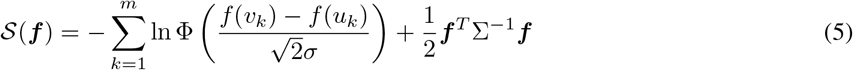

With ***f***_MAP_ obtained, the evidence can then be approximated as

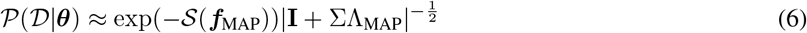

The approximation in 6 can be solved analytically and is differentiable, allowing for hyperparameter optimization through gradient descent.

We optimize PKermut using the AdamW optimizer with a learning rate of 0.1 for 150 steps, identical to Kermut’s training routine. We perform training and inference at float64 precision to increase numerical stability, as per the recommendations of botorch. We obtain per-sample posterior predictions directly from the learned latent function after optimizing PKermut hyperparameters.

**Augmented Potts** is a ridge regression model with inputs comprised of flattened one-hot sequence encodings and a sequence energy score assigned by a Potts model. Sequence energy scores are derived from multiple sequence alignments (MSAs) provided by the ProteinGym, c.f. [17].

**RITA regressor** is like Augmented Potts, a ridge regression model, but instead using RITA_xl embeddings mean-pooled over the sequence length, c.f. [17].

**Training** All models are trained on all datasets using 5-fold quantile CV. Spearman scores on test folds are recorded, and posterior predictions for the test data for each training run are stored for later acquisition experiments.

### 4.4 Acquisition functions

### 4.5 Glossary

*x*_*i*_ Variant with sample index *i*.

*y*_*i*_ Ground truth property value of *x*_*i*_.

*y*_*ref*_ Ground truth property value of a reference variant; in our analysis, this is the variant with the highest property value in the training set.

*I*(*x*_*i*_) The improvement of *y*_*i*_ over *y*_*ref*_: *I*(*x*_*i*_) = max(*y*_*ref*_ *− y*_*i*_, 0).

*µ*_*i*_ The posterior mean of *x*_*i*_.

*σ*_*i*_ The posterior standard deviation of *x*_*i*_.

Φ The cumulative density function of a standard Gaussian distribution.

*ϕ* The probability density function of a standard Gaussian distribution.

*a*_*i*_ The acquisition value of *x*_*i*_ obtained by any of the acquisition functions.

The posterior distributions derived from the models are assumed to be Gaussian, such that the posterior prediction of *x*_*i*_ can be denoted by 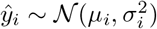.

#### 4.5.1 Definitions

##### Greedy

Uncertainty unaware acquisition that ranks variants by their predicted posterior mean, *a*_Greedy,*i*_ = *µ*_*i*_.

##### Probability of improvement

Uncertainty-aware acquisition that selects variants by the predicted posterior probability that *I*(*x*_*i*_) is greater than 0. Acquisition values are obtained as follows:

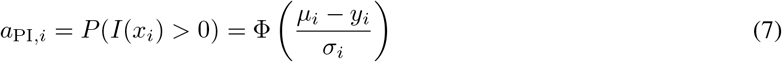

As *σ*_*i*_ increases, corresponding to higher predictive uncertainty, *a*_*i*_ decreases.

##### Expected improvement

Uncertainty-aware acquisition that ranks variants by the posterior expectation over *I*(*x*_*i*_). Acquisition values are obtained by:

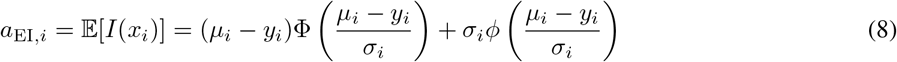

Whereas the probability of improvement only considers the probability that *I*(*x*_*i*_) is positive, the expected improvement puts more emphasis on the magnitude of improvement. The second term ensures that *a*_EI,*i*_ increases with increased values of *σ*_*i*_.

##### Thompson sampling

Uncertainty aware acquisition that ranks variants by the values obtained by drawing a random sample from their predicted posteriors, 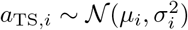.

##### Upper confidence bound

Uncertainty aware acquisition that ranks variants by the upper confidence bound of their predicted posterior, which is calculated as *a*_UCB,*i*_ = *µ*_*i*_ +*βσ*_*i*_. Increased values of *β* correspond to increased uncertainty reward. In all our acquisition experiments, we fix *β* = 0.1.

### 4.6 Critical differences diagram

We plot the CD diagram using the scikit-posthocs library [24]. The scikit-posthocs library determines statistical significance of differences by computing Conover post-hoc tests for each pair of strategies.

## Data availability

We obtained all datasets used in this analysis from the ProteinGym [9].

## Code availability

Code is made available at https://github.com/florisvdf/kermut-package.

## Declaration of generative AI and AI-assisted technologies in the writing process

During the writing of the article, the authors used various generative models developed by OpenAI and Anthropic to assist with grammar, flow, and overall refinement of the text. After using these tools, the authors carefully reviewed and edited the generated content as needed and take full responsibility for the content of the published article.

## A Supplementary Materials

### A.1 Excluded selection assay categories

The following is a list of categories under the “selection_assay” column in the ProteinGym metadata for which no dataset could be found that met our inclusion criteria and that we could train all our models on without exceeding our computational budget.

“viral replication”, “uptake of cytotoxic substrate”, “Binding to Mcl-1 (FACS; yeast-displayed and antibody stained for binding partner)”, “aggregation”, “Viral Replication”, “Expression”, “Drug resistance”, “viability for AAV capsid production”, “growth, nitrogen depletion (0.0125% ammonium sulfate), hyperosmotic shock (0.8 M NaCl), alcohol stress (7.5% ethanol), sulfhydryl-oxidation (0.85 mM diamide), temperature shock (37C)”, “Fitness”, “nan”, “growth (surrogate for toxicity/activity of CCDB)”, “Viral replication (avian cells: CCL141 (duck))”, “Voltage”, “growth/survival (surrogate for TpoR/MPL enhanced constitutive activation)”, “drug resistance (surrogate for protein activity, 6-thioguanine (6-TG))”, “Growth (no selection, 24h)”, “Binding”, “drug resistance (triple-drug assay: veratridine + brevetoxin + ouabain; surrogate for sodium channel dysfunction, select against function)”, “peptide binding”, “Ligase activity (phage display)”, “Membrane-protein insertion”.

**Table S1:**
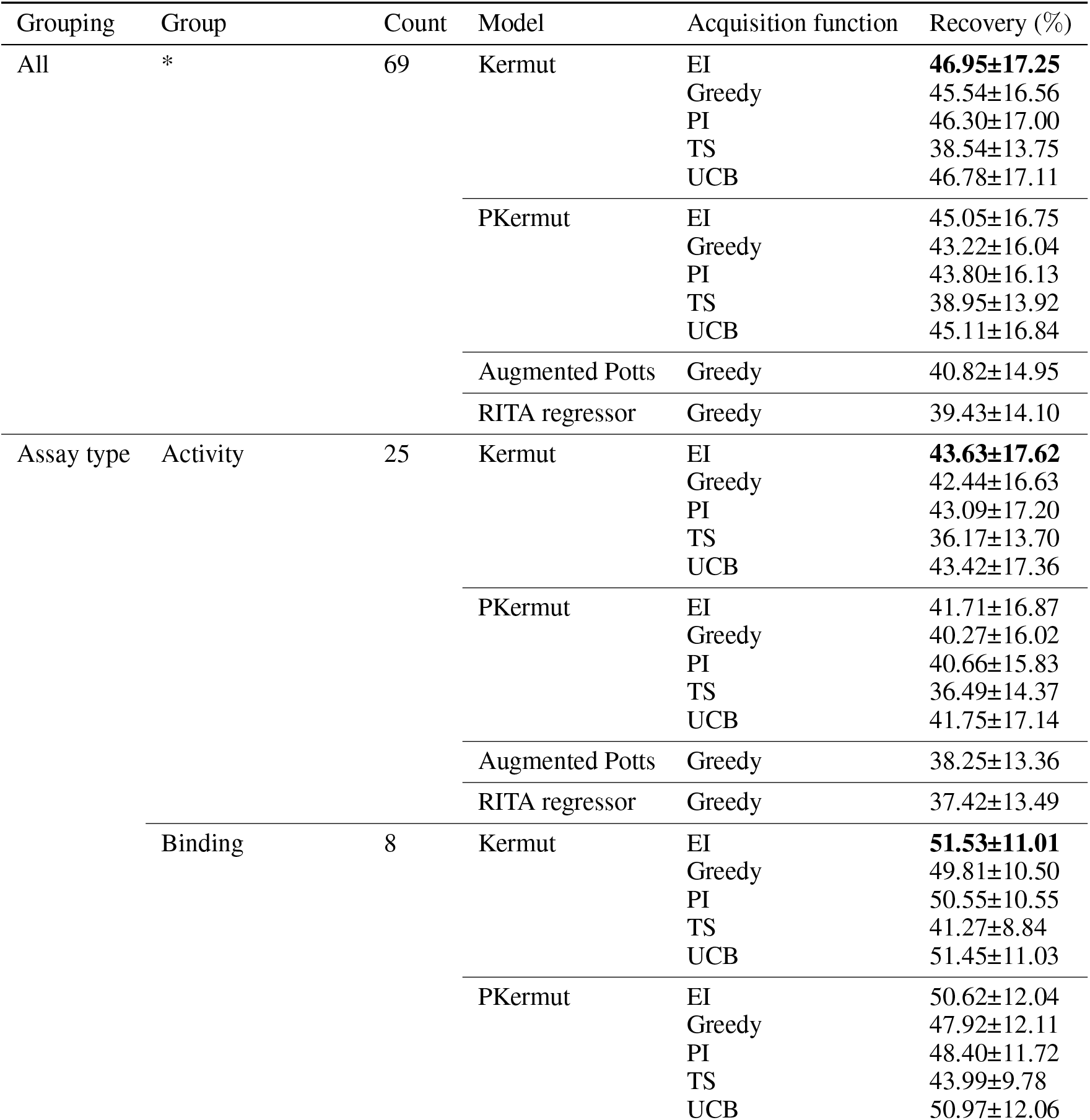
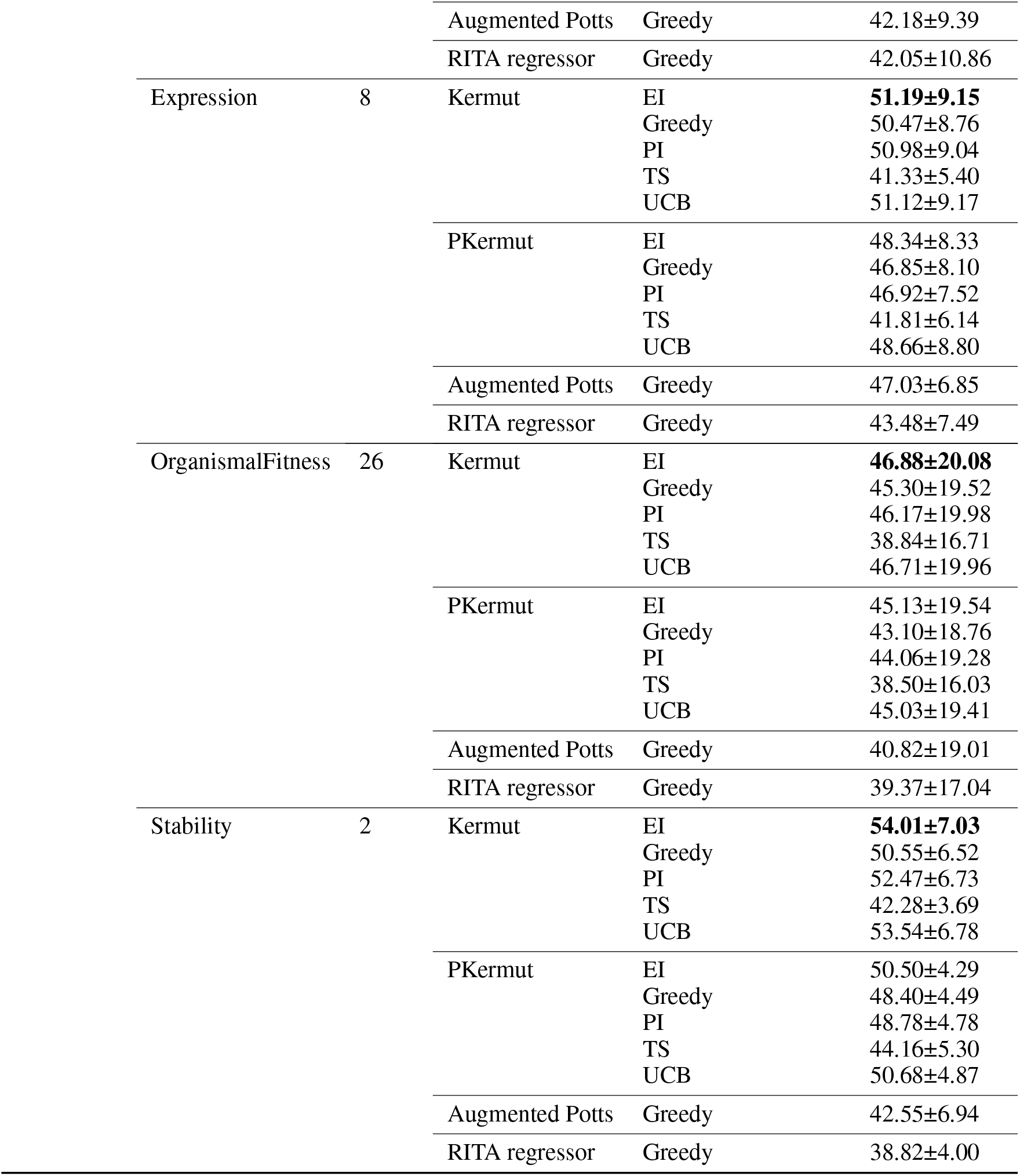
Acquisition recovery scores for every combination of dataset group, model, and acquisition function.

**Figure S1:**
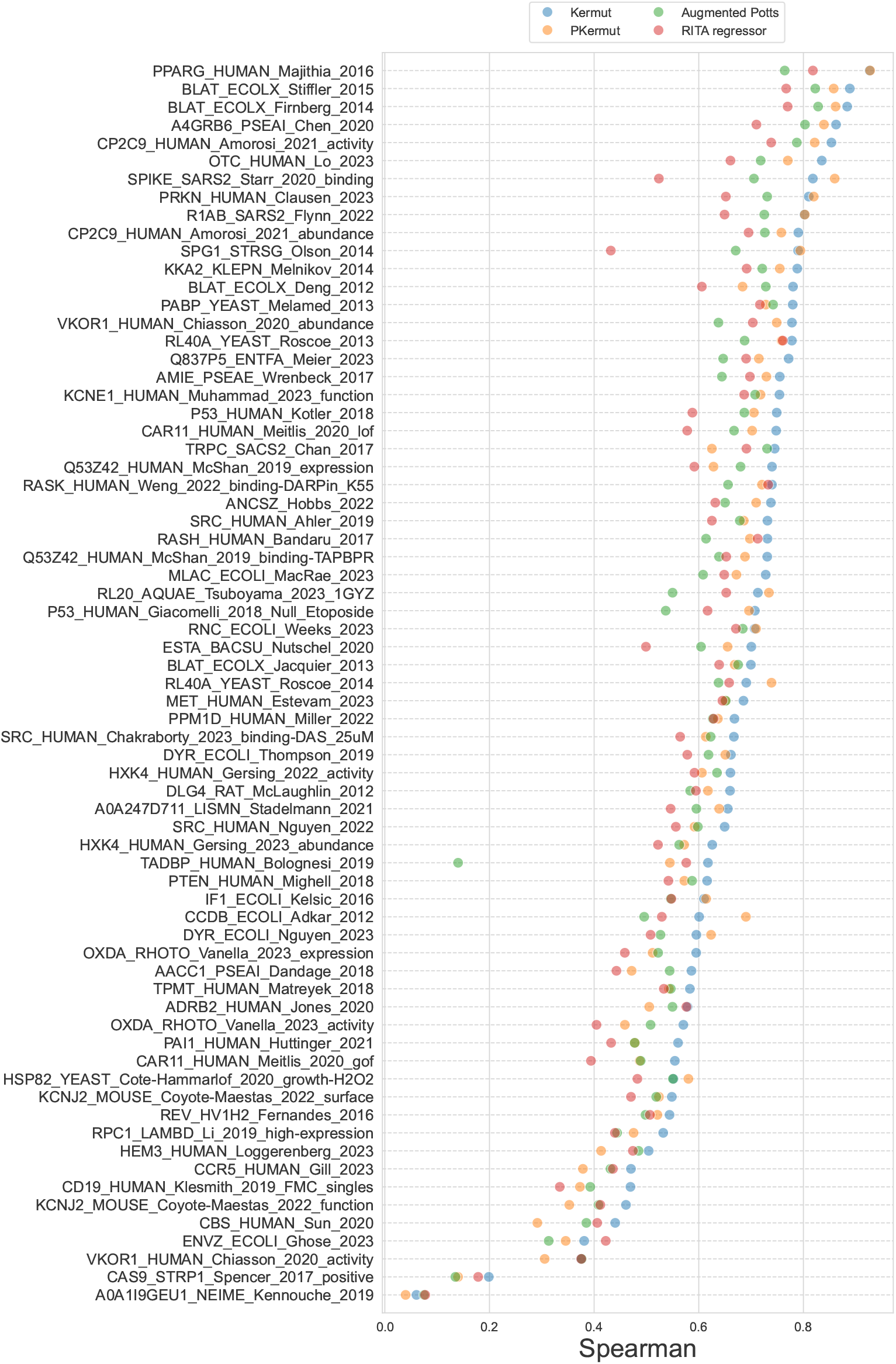
Test-fold averaged Spearman correlation scores of all models on each dataset using the quantile CV strategy, ordered by the scores of Kermut.

## References

[1] Christian M Heckmann and Francesca Paradisi. Looking back: a short history of the discovery of enzymes and how they became powerful chemical tools. ChemCatChem, 12(24):6082–6102, 2020.

[2] Frances H Arnold. Protein engineering for unusual environments. Current opinion in biotechnology, 4(4):450–455, 1993.

[3] Brian L Hie and Kevin K Yang. Adaptive machine learning for protein engineering. Current opinion in structural biology, 72:145–152, 2022.

[4] Chase R Freschlin, Sarah A Fahlberg, and Philip A Romero. Machine learning to navigate fitness landscapes for protein engineering. Current opinion in biotechnology, 75:102713, 2022.

[5] Petr Kouba, Pavel Kohout, Faraneh Haddadi, Anton Bushuiev, Raman Samusevich, Jiri Sedlar, Jiri Damborsky, Tomas Pluskal, Josef Sivic, and Stanislav Mazurenko. Machine learning-guided protein engineering. ACS catalysis, 13(21):13863–13895, 2023.

[6] Jason Yang, Ravi G Lal, James C Bowden, Raul Astudillo, Mikhail A Hameedi, Sukhvinder Kaur, Matthew Hill, Yisong Yue, and Frances H Arnold. Active learning-assisted directed evolution. Nature Communications, 16(1): 714, 2025.

[7] Tudor-Stefan Codet and Igor Krawczuk. Protein optimization 101: Insights from the literature, Oct 2024. URL https://www.adaptyvbio.com/blog/po101/.

[8] Peter Mørch Groth, Mads Kerrn, Lars Olsen, Jesper Salomon, and Wouter Boomsma. Kermut: Composite kernel regression for protein variant effects. Advances in Neural Information Processing Systems, 37:29514–29565, 2024.

[9] Pascal Notin, Aaron Kollasch, Daniel Ritter, Lood Van Niekerk, Steffanie Paul, Han Spinner, Nathan Rollins, Ada Shaw, Rose Orenbuch, Ruben Weitzman, et al. Proteingym: Large-scale benchmarks for protein fitness prediction and design. Advances in Neural Information Processing Systems, 36, 2024.

[10] Alexander D’Amour, Katherine Heller, Dan Moldovan, Ben Adlam, Babak Alipanahi, Alex Beutel, Christina Chen, Jonathan Deaton, Jacob Eisenstein, Matthew D Hoffman, et al. Underspecification presents challenges for credibility in modern machine learning. Journal of Machine Learning Research, 23(226):1–61, 2022.

[11] Brian Hie, Bryan D Bryson, and Bonnie Berger. Leveraging uncertainty in machine learning accelerates biological discovery and design. Cell systems, 11(5):461–477, 2020.

[12] Kevin P Greenman, Ava P Amini, and Kevin K Yang. Benchmarking uncertainty quantification for protein engineering. PLOS Computational Biology, 21(1):e1012639, 2025.

[13] FLIP: Benchmark tasks in fitness landscape inference for proteins, author=Dallago, Christian and Mou, Jody and Johnston, Kadina E and Wittmann, Bruce J and Bhattacharya, Nicholas and Goldman, Samuel and Madani, Ali and Yang, Kevin K. bioRxiv, pages 2021–11, 2021.

[14] Kieran Didi, Sarah Alamdari, Alex X Lu, Bruce Wittmann, Kadina E Johnston, Ava P Amini, Ali K Madani, Maya Czeneszew, Christian Dallago, and Kevin K Yang. FLIP2: Expanding Protein Fitness Landscape Benchmarks for Real-World Machine Learning Applications. bioRxiv, 2026.

[15] Matthew Hoffman, Eric Brochu, Nando De Freitas, et al. Portfolio Allocation for Bayesian Optimization. In UAI, volume 11, pages 327–336, 2011.

[16] Wei Chu and Zoubin Ghahramani. Preference learning with Gaussian processes. In Proceedings of the 22nd international conference on Machine learning, pages 137–144, 2005.

[17] Floris van der Flier, Dave Estell, Sina Pricelius, Lydia Dankmeyer, Sander van Stigt Thans, Harm Mulder, Rei Otsuka, Frits Goedegebuur, Laurens Lammerts, Diego Staphorst, et al. Enzyme structure correlates with variant effect predictability. Computational and Structural Biotechnology Journal, 23:3489–3497, 2024.

[18] Eli Bixby, Gino Brunner, Daniel Danciu, Richard Dela Rosa, Nicolas Deutschmann, Constance Ferragu, Franziska Geiger, Christian Holberg, Patrick Kidger, Arthur Lindoulsi, Noé Lutz Thomas McColgan, Sebastian Milius, Jinel Shah, Michelle Vandeloo, Paula Vidas, Jonathan D. Ziegler, Harmen van Rossum, Daan van der Vorm, Nicolò Baldi Catalina IJSpeert, Emanuele Monza, and Angela Schriek. What comes after de novo? Automated lead optimization of proteins with CRADLE-1. bioRxiv, 2026.

[19] William BT Mock. Pareto optimality. In Encyclopedia of global justice, pages 808–809. Springer, 2011.

[20] Ava P Soleimany, Alexander Amini, Samuel Goldman, Daniela Rus, Sangeeta N Bhatia, and Connor W Coley. Evidential deep learning for guided molecular property prediction and discovery. ACS central science, 7(8): 1356–1367, 2021.

[21] Zeming Lin, Halil Akin, Roshan Rao, Brian Hie, Zhongkai Zhu, Wenting Lu, Nikita Smetanin, Robert Verkuil, Ori Kabeli, Yaniv Shmueli, et al. Evolutionary-scale prediction of atomic-level protein structure with a language model. Science, 379(6637):1123–1130, 2023.

[22] Justas Dauparas, Ivan Anishchenko, Nathaniel Bennett, Hua Bai, Robert J Ragotte, Lukas F Milles, Basile IM Wicky, Alexis Courbet, Rob J de Haas, Neville Bethel, et al. Robust deep learning–based protein sequence design using ProteinMPNN. Science, 378(6615):49–56, 2022.

[23] Maximilian Balandat, Brian Karrer, Daniel Jiang, Samuel Daulton, Ben Letham, Andrew G Wilson, and Eytan Bakshy. BoTorch: A framework for efficient Monte-Carlo Bayesian optimization. Advances in neural information processing systems, 33:21524–21538, 2020.

[24] Maksim A Terpilowski. scikit-posthocs: Pairwise multiple comparison tests in Python. Journal of Open Source Software, 4(36):1169, 2019.

